# *Plasmodium berghei* liver stages establish a damage-mimicking vacuole to hijack host ER membrane contact site machinery for lipid uptake

**DOI:** 10.64898/2026.06.30.735545

**Authors:** Dominik Liebeck, Jacqueline Schmuckli-Maurer, Stephan Hirschi, Michaela Bulloch, Reto Caldelari, Nina Urban, Virginia Grünig, Aymeric Dossmann Ritter, Alva Josefin Assmann, Philipp Olias, Volker T. Heussler, Paul-Christian Burda

**Affiliations:** Institute of Veterinary Pathology, Justus Liebig University Giessen, 35392 Giessen, Germany; Institute of Cell Biology, University of Bern, 3012 Bern, Switzerland; Institute of Biochemistry and Molecular Medicine, University of Bern, 3012 Bern, Switzerland

## Abstract

*Plasmodium* liver stage parasites undergo massive intracellular replication and therefore require extensive acquisition of host-derived lipids. How parasites coordinate lipid supply at the host-parasite interface, however, remains poorly understood. Here, we show that the *Plasmodium berghei* parasitophorous vacuole membrane (PVM) selectively mimics features of damaged endolysosomal membranes, including dynamic LC3-positive membrane tubules and PI4K2A-dependent PI(4)P accumulation, while lacking canonical damage markers including ESCRT components and galectins. This damage-mimicking membrane state drives OSBP recruitment and is associated with formation of VAPA/B-positive ER-PVM membrane contact sites. Disruption of VAPA/B or OSBP impaired PVM integrity and parasite survival, while OSBP depletion reduced cholesterol accumulation at the PVM. Together, our findings suggest that liver stage parasites exploit host lysosome repair-like pathways to redirect host lipids toward parasite growth and intracellular survival. These results redefine the PVM as an active metabolic interface that enables host lipid acquisition rather than merely serving as a protective barrier.

## INTRODUCTION

*Plasmodium* liver stage parasites undergo massive intracellular replication and therefore require extensive acquisition of host-derived lipids. A single invading sporozoite differentiates into a multinucleated schizont that produces thousands of merozoites within days, necessitating substantial membrane biogenesis. While *Plasmodium* encodes enzymes for *de novo* lipid synthesis^1^, liver stages rely heavily on host-derived lipids^2–5^. In particular, *Plasmodium* lacks sterol biosynthesis pathways and is therefore entirely dependent on host-derived cholesterol^6^. How liver stage parasites orchestrate lipid acquisition at the host-parasite interface remains incompletely understood.

Upon hepatocyte invasion, *Plasmodium* resides within a parasitophorous vacuole (PV) surrounded by the PV membrane (PVM). Beyond serving as a protective barrier, the PVM acts as a dynamic host-parasite interface that recruits host proteins involved in vesicular trafficking, autophagy-related pathways, and metabolic adaptation^7–10^. One striking feature of the liver stage PVM is its persistent association with host LC3^11–13^. Unlike canonical autophagy, LC3 recruitment is stable and does not culminate in parasite elimination^13^. Instead, LC3 lipidation depends on the V-ATPase-ATG16L1 axis^9^, placing this response within the framework of conjugation of ATG8 to single membranes (CASM). CASM is increasingly recognized as a response to perturbed endolysosomal membranes that promotes membrane remodeling and restoration of homeostasis^14,15^. The sustained engagement of CASM machinery at the PVM therefore raises the possibility that *Plasmodium* establishes a membrane state resembling damaged endolysosomal compartments.

Damaged lysosomes engage a coordinated membrane repair response characterized by recruitment of LC3, phosphoinositide remodeling, and formation of membrane contact sites (MCSs) with the endoplasmic reticulum (ER)^16–18^. Key components of this response include PI4K2A-dependent generation of phosphatidylinositol 4-phosphate (PI(4)P) at the damaged membrane, recruitment of OSBP/ORP-family lipid transfer proteins, and formation of VAPA/B-dependent ER MCSs that facilitate lipid transport and membrane restoration.

Here, we show that *Plasmodium berghei* liver stages establish a PVM that mimics key features of damaged endolysosomal membranes. The PVM lacks damage markers such as ESCRT components and galectins, yet displays PI4K2A-dependent PI(4)P accumulation, forms VAPA/B-positive ER-PVM MCSs, and recruits OSBP. Disruption of VAPA/B or OSBP compromises PVM integrity and parasite survival, while OSBP depletion additionally reduces cholesterol accumulation at the host-parasite interface. Together, our findings identify damage mimicry as a previously unrecognized strategy by which *Plasmodium* co-opts host membrane remodeling and lipid transfer pathways.

## RESULTS

### The liver stage PVM forms dynamic LC3-positive tubules resembling structures associated with lysosomal membrane damage

To identify early host cell responses associated with the *P. berghei* liver stage PVM, we performed live-cell imaging of infected host cells expressing fluorescently tagged LC3. Strikingly, we observed highly dynamic LC3-positive membrane tubules emanating from the PVM of early liver stage parasites (Figure 1A, Supplementary Movie S1), which frequently extended several micrometers into the host cytosol and underwent rapid extension, retraction and remodeling. Morphologically and dynamically, these structures resembled LC3-positive tubules recently described at damaged lysosomes during CASM responses^15^. Like these lysosomal tubules, PVM tubule formation required intact microtubules, as nocodazole treatment reduced the proportion of tubule-positive parasites from 62% to 23% (Figure 1B). PVM tubule formation was also observed in primary hepatocytes derived from transgenic mice ubiquitously expressing GFP-LC3^19^ (Figure 1C). To determine whether this membrane state is established during physiological infection, we performed intravital microscopy using GFP-LC3 expressing mice infected with fluorescent *P. berghei* parasites. LC3-positive PVM tubules were also detected in living mice (Figure 1D, Supplementary Movie S2), confirming the *in vivo* relevance of this response. These observations prompted us to examine additional hallmarks of lysosomal membrane stress active at the PVM.

**Figure 1.**
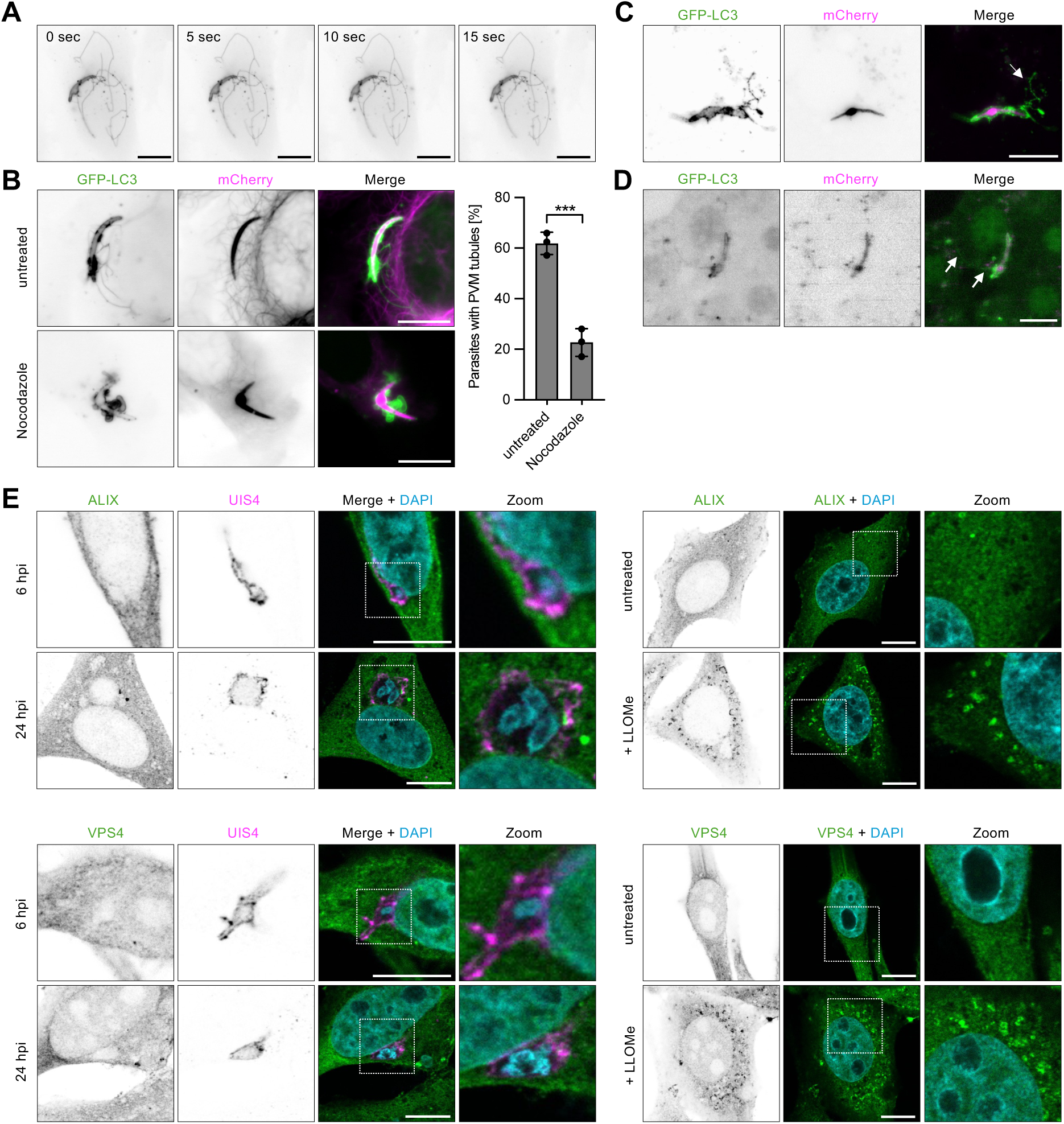
The liver stage PVM forms LC3-positive membrane tubules characteristic of damaged lysosomes *in vitro* and *in vivo*, but does not recruit host ESCRT components. (A) Stills from a live-cell movie of *P. berghei*-infected HeLa cell stably expressing GFP-LC3, showing dynamic LC3-positive membrane tubules at the PVM. Movie started at 4 hours post-infection (hpi). See also Movie S1. (B) Quantification of LC3-positive PVM tubule-forming parasites at 4 hpi in HeLa cells expressing the microtubule binding-domain of ensconsin fused to mCherry (EMTB-mCherry), which is visualized in the same channel as the cytoplasm of mCherry expressing parasites (magenta). Infected cells were left untreated or treated for 150 min with 20 µM nocodazole. Shown are means ± SD from 3 independent experiments. Statistical significance was assessed by Welch’s t-test. ***p < 0.001. Representative images of an untreated and nocodazole-treated parasite are shown on the left. (C) LC3-positive PVM tubules are also visible in infected primary mouse hepatocytes isolated from GFP-LC3 mice at 4 hpi. PVM tubules are highlighted by arrow. (D) Still from an intravital microscopy movie of a *P. berghei*-infected GFP-LC3 mouse at 4 hpi, showing LC3-positive membrane tubules at the liver stage PVM *in vivo*. PVM tubules are highlighted by arrows. See also Movie S2. (E) IFA shows that neither ESCRT component ALIX nor the ESCRT-associated AAA ATPase VPS4 localizes to the PVM at 6 and 24 hpi. The PVM was visualized by staining against the PVM marker protein UIS4. As a positive control, lysosomal damage was induced by treating cells with 250 µM LLOMe for one hour. Scale bars, 10 µm.

### The liver stage PVM lacks recruitment of host ESCRT membrane repair machinery

To determine whether the liver stage PVM engages membrane repair pathways associated with lysosomal membrane damage, we next examined recruitment of the ESCRT machinery component ALIX and the ESCRT-associated AAA ATPase VPS4. In mammalian cells, damaged lysosomes rapidly recruit ESCRT factors including ALIX that contribute to membrane repair following membrane injury^20,21^. In contrast to the pronounced LC3-positive tubulation observed at the PVM, neither ALIX-mNeonGreen nor GFP-VPS4 accumulated at the PVM of *P. berghei* liver stages (Figure 1E). As a positive control, treatment of uninfected cells with the lysosomotropic compound LLOMe induced ALIX- and VPS4-positive foci, consistent with efficient recruitment of the ESCRT membrane repair machinery to damaged lysosomes (Figure 1E). These findings are consistent with previous observations that the liver stage PVM is not recognized by cytosolic galectins^13^, which detect cytosolically exposed luminal glycans following lysosomal rupture^22^. Together, these observations suggest that the PVM shares selected features with stressed or damaged lysosomal membranes while lacking both ESCRT-associated membrane repair responses and galectin-mediated recognition of membrane rupture.

### Host PI4K2A generates a PI(4)P-positive membrane state at the liver stage PVM

To determine whether the liver stage PVM activates phosphoinositide signaling pathways associated with damaged lysosomes, we first examined localization of PI4K2A, a kinase that generates PI(4)P during lysosomal membrane stress responses^16,17^. Strikingly, PI4K2A-HA was clearly recruited to the PVM as early as 2 hours post-infection (hpi), as visualized by co-localization with the PVM marker UIS4, and remained associated with the PVM throughout liver stage development (Figure 2A). PI4K2A recruitment occurred independently of ATG16L1, indicating independence of CASM (Figure 2A). Next, the localization of phosphoinositides was analyzed using fluorescent lipid reporters in live infected cells. Among these, only the PI(4)P reporter displayed robust enrichment around the parasite (Figure 2B). Immunofluorescence analysis (IFA) confirmed co-localization of the PI(4)P reporter with UIS4 (Figure 2C), indicating that PI(4)P is generated at the PVM. Complementing this observation, the liver stage PVM was previously shown to be enriched in phosphatidylserine^9^, a lipid that, together with PI(4)P, is characteristic of damaged lysosomal membranes^16,17^. To determine whether the PI(4)P enrichment at the PVM depends on host PI4K2A, we analyzed PI(4)P localization in PI4K2A-knockout (KO) cells^16^. Loss of PI4K2A significantly reduced both the proportion of PI(4)P-positive parasites and relative PI(4)P reporter signal intensity at the PVM (Figure 2C). PI(4)P accumulation was restored by PI4K2A complementation. Unexpectedly, PI4K2A-deficient host cells displayed a modest increase in parasite size and survival that was reverted upon PI4K2A complementation (Figure S1A,B). These findings indicate that PI4K2A-dependent PI(4)P production is not required for parasite survival and suggest that host PI4K2A signaling at the PVM may restrict liver stage development under the conditions tested.

**Figure 2.**
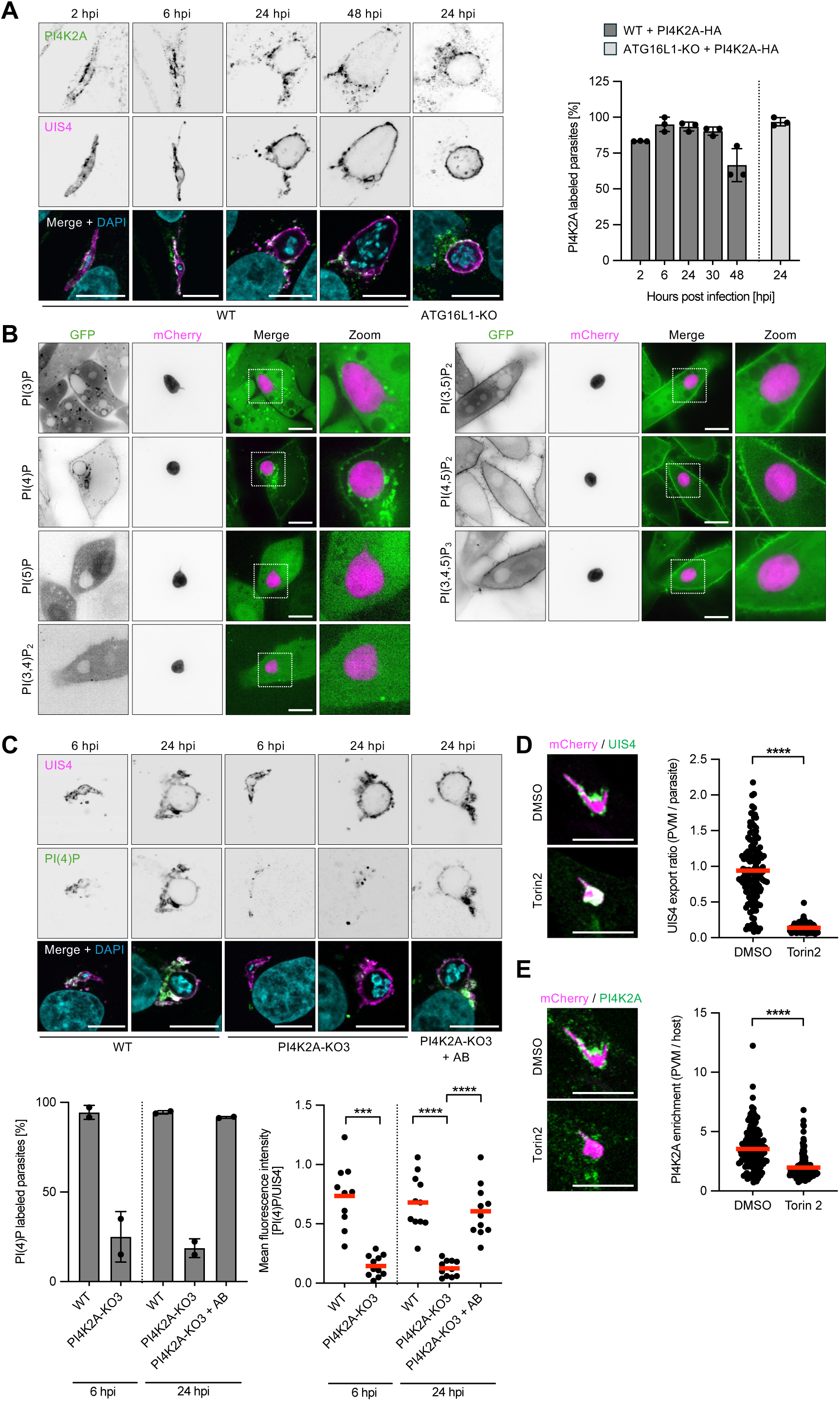
Host PI4K2A is recruited to the PVM and contributes to PI(4)P production at this membrane. (A) IFA of infected HeLa cells reveals that PI4K2A-HA is recruited to the PVM throughout liver stage development independent of CASM. For the quantifications, means ± SD of three independent experiments are shown. WT, wildtype; KO, knockout. (B) Live-cell microscopy of fluorescent phosphoinositide reporters in infected HeLa cells at 24-29 hpi reveals that liver stage parasites are associated with PI(4)P. (C) The PI(4)P reporter co-localizes with the PVM marker UIS4 in IFA. PI(4)P levels at the PVM are significantly reduced in PI4K2A-KO cells and this effect is rescued by PI4K2A complementation. For the percentage of PI(4)P-labelled parasites means ± SD are shown, while for the mean fluorescence intensity analysis of the PI(4)P reporter, data points derived from individual parasites with means indicated in red are displayed. Data are derived from two independent experiments. Statistical significance was assessed by Welch’s ANOVA followed by Dunnett’s T3 multiple-comparison test. ***p < 0.001, ****p < 0.0001. AB, PI4K2A-HA addback. See also Figure S1. (D, E) 10 nM Torin2 treatment from 2 to 8 hpi blocks export of UIS4 to the PVM and impairs recruitment of host PI4K2A to the PVM. Shown are pooled data of four independent experiments with means indicated in red. Statistical significance was assessed by Welch’s t-test. ****p < 0.0001. See also Figure S2. Scale bars, 10 µm.

To examine how PI4K2A becomes enriched at the PVM, we treated infected cells with Torin2, which inhibits parasite protein export to the PVM^23^. As expected, Torin2 efficiently blocked export of UIS4 (Figure 2D, Figure S2). Remarkably, Torin2 treatment also strongly reduced PI4K2A recruitment to the PVM (Figure 2E, Figure S2), suggesting that one or more parasite-exported proteins are required for its accumulation at this membrane.

### Host VAPA and VAPB localize to ER-PVM contact sites and are required for successful parasite development and proper PVM morphology

At damaged lysosomes, PI(4)P accumulation drives ER recruitment and formation of ER-lysosome MCSs via the ER tethering proteins VAPA and VAPB^16,17^. Given that the liver stage PVM similarly accumulates PI(4)P, and that the host ER has been shown to locally accumulate around the liver stage PV^24^, we next asked whether the PVM recruits VAPA and VAPB to establish analogous ER-PVM MCSs. IFA of liver stage-infected host cells expressing GFP-VAPA or GFP-VAPB revealed their partial co-localization with the PVM marker UIS4 (Figure 3A). To further test whether VAP-positive ER membranes closely associate with the PVM, we performed proximity ligation assays (PLAs), which generate a signal only when two epitopes are located within ∼40 nm of each other, and are widely used to assess MCS formation^25^. For these experiments, VAPA/B double knockout (VAP-DKO) cells^26^ stably expressing HA-VAPB were infected with parasites and analyzed by PLA using antibodies against HA and UIS4. PVM-associated PLA puncta were readily detected in HA-VAPB-expressing cells, whereas essentially no PLA puncta were detected in infected wildtype (WT) cells lacking HA-tagged VAPB (Figure 3B), confirming assay specificity. These findings are consistent with formation of VAPA/B-positive ER-PVM MCSs during liver stage infection.

**Figure 3.**
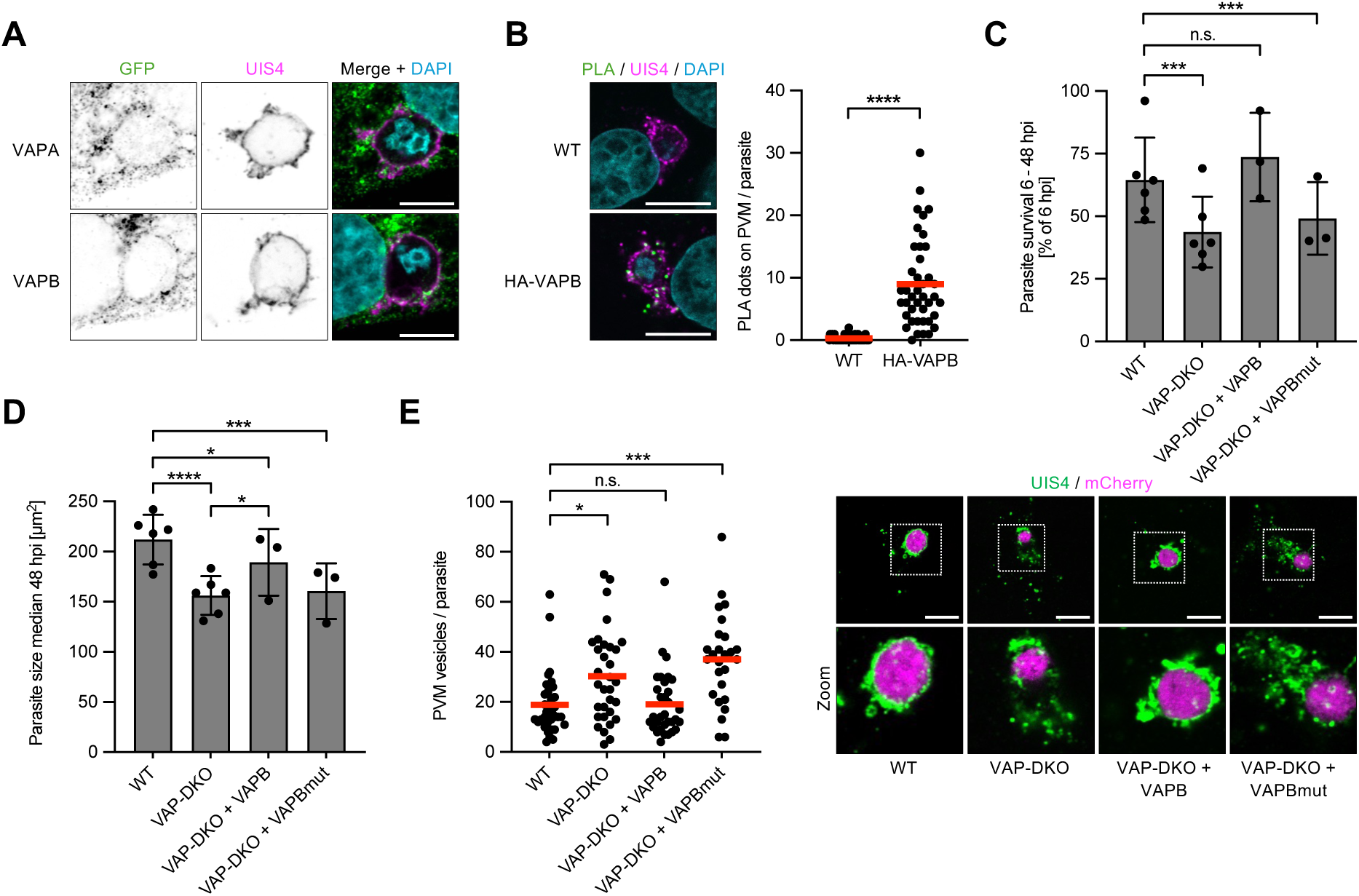
Host ER-tethering proteins VAPA and VAPB are found at ER-PVM MCSs and contribute to parasite development and PVM integrity through their FFAT motif-binding activity. (A) IFA of infected HeLa cells shows that GFP-VAPA and GFP-VAPB partially co-localize with the PVM marker UIS4. (B) PLA of WT and HA-VAPB-expressing infected cells at 24 hpi using antibodies against HA and UIS4 reveals VAPB at ER-PVM MCSs. Shown are pooled data of two independent experiments with means indicated in red. Statistical significance was assessed by Welch’s t-test. ****p < 0.0001. (C,D) Parasite survival from 6 to 48 hpi (C) and size at 48 hpi (D) are decreased in VAP-DKO cells. Complementation with WT VAPB restores these phenotypes, while mutated VAPB (VAPBmut) unable to interact with FFAT-motifs does not complement. Shown are means ± SD of three to six independent experiments. Statistical significance was assessed by mixed-effects model (REML) with experiment treated as a matched factor, followed by Holm–Šidák multiple-comparison testing. *p < 0.05, ***p < 0.001, ****p < 0.0001; n.s., not significant. (E) IFA reveals that PVM integrity is impaired in VAP-DKO cells as visible by hypervesiculation. Complementation with WT VAPB rescues this phenotype, while VAPBmut does not. Shown are pooled data of two independent experiments with means indicated in red. Statistical significance was assessed by Welch’s ANOVA followed by Dunnett’s T3 multiple-comparison test. *p < 0.05, ***p < 0.001; n.s., not significant. Representative IFA images of the PVM morphology in the different cell lines are shown. Scale bars,10 µm.

To determine the functional importance of VAPA/B during liver stage development, we compared parasite development in WT and VAP-DKO cells. Parasite survival and parasite size at 48 hpi were significantly reduced in VAP-DKO cells (Figure 3C,D). Both phenotypes were restored by complementation with WT VAPB but not with a mutant variant of VAPB defective in interactions with FFAT motif-containing proteins (K87D/M89D, VAPBmut), indicating that VAPA/B-mediated membrane tethering contributes to parasite development. In addition to impaired parasite growth and survival, loss of VAPA/B strongly affected PVM morphology. Parasites developing in VAP-DKO cells displayed pronounced hypervesiculation of the PVM, as quantified by automated image analysis (Figure 3E). The pronounced alterations in PVM morphology precluded reliable quantitative analysis of ER-PVM association in VAP-DKO cells. PVM hypervesiculation was rescued by re-expression of WT VAPB but not VAPBmut, further supporting an important role for VAPA/B in maintenance of PVM integrity.

### Host OSBP is recruited to the PVM in a PI4K2A-dependent manner and contributes to PVM cholesterol accumulation and integrity

OSBP is recruited to PI(4)P-positive membranes through its PI(4)P-binding PH domain and has been identified as a component of the membrane repair response at damaged lysosomes, where it interacts with VAPA/B at ER-lysosome MCSs^16,17^. We therefore asked whether the PI(4)P-enriched liver stage PVM similarly recruits OSBP. In WT cells expressing GFP-OSBP, OSBP partially co-localized with UIS4. Notably, OSBP recruitment to the PVM was strongly reduced in PI4K2A-KO cells and restored upon complementation with PI4K2A-HA. Quantification of Pearson’s correlation coefficients between UIS4 and GFP-OSBP confirmed significantly reduced co-localization in PI4K2A-KO cells compared to WT and complemented cells (Figure 4A). These findings indicate that recruitment of OSBP to the PVM depends on host PI4K2A and are consistent with a model in which PI4K2A-generated PI(4)P at the PVM promotes recruitment of OSBP to this membrane.

**Figure 4.**
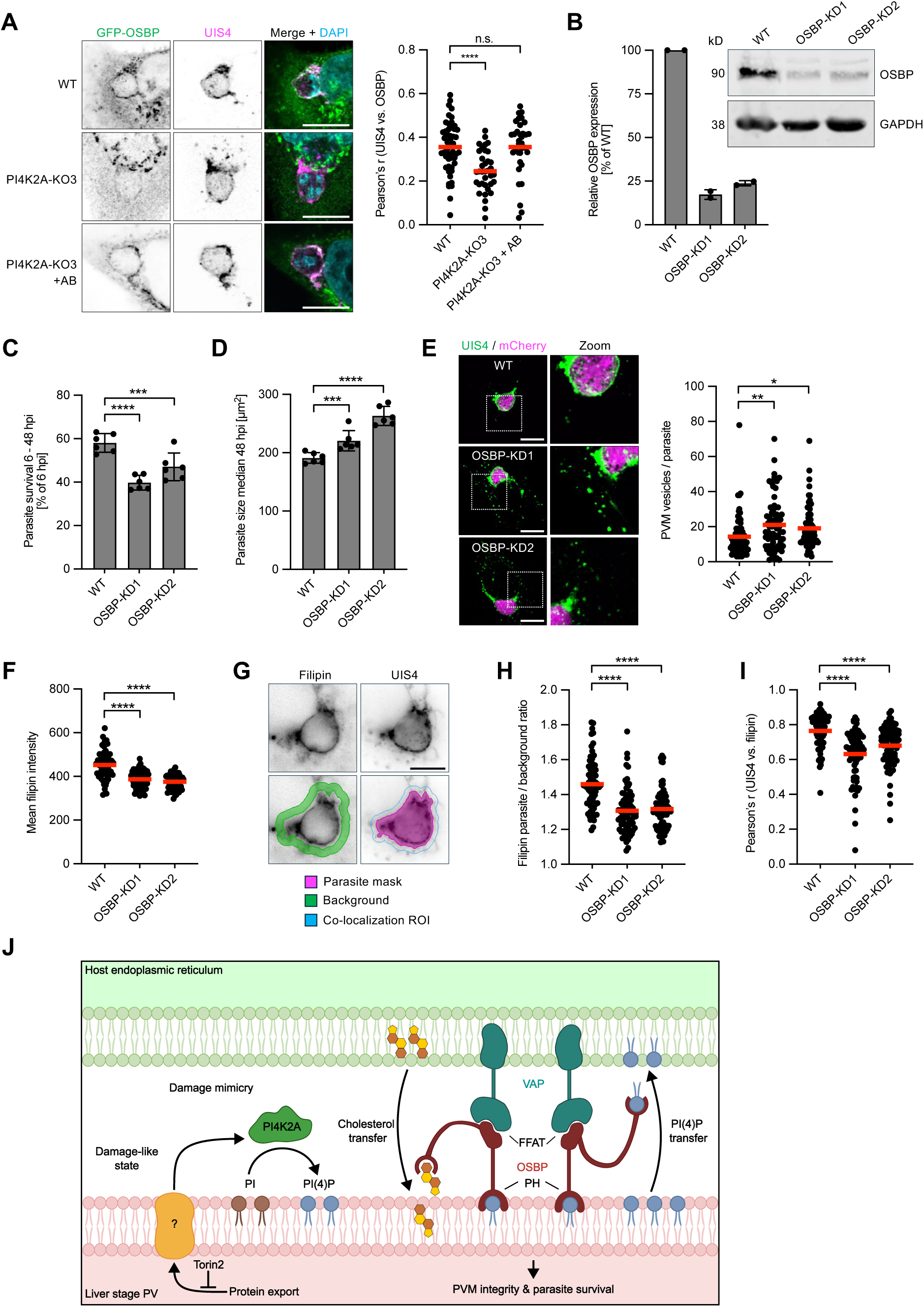
Host OSBP localizes to the PVM in a PI4K2A-dependent manner and is important for PVM integrity and cholesterol accumulation at this membrane. (A) Co-localization analysis of GFP-OSBP and the PVM marker UIS4 in WT, PI4K2A-KO and complemented PI4K2A-KO cells by IFA reveals PI4K2A-dependent OSBP recruitment to the PVM. For the quantification of the Pearsons correlation coefficient between GFP-OSBP and UIS4, pooled data of two to three independent experiments are shown with means indicated in red. Statistical significance was assessed by Welch’s ANOVA followed by Dunnett’s T3 multiple-comparison test. ****p < 0.0001; n.s., not significant. AB, PI4K2A-HA addback. (B) Verification of OSBP-KD by Western blot. OSBP expression was normalized to GAPDH with WT cells set to 100% in each experiment. Shown are means ± SD of two independent experiments. (C) Parasite survival (6 to 48 hpi) is decreased in OSBP-KD cells. (D) Parasite size at 48 hpi is increased in OSBP-KD cells. For (C) and (D) means ± SD of six independent experiments are shown. Statistical significance was assessed by repeated-measures one-way ANOVA followed by Holm–Šidák multiple-comparison testing. ***p < 0.001, ****p < 0.0001. (E) IFA reveals impaired PVM integrity in OSBP-KD cells. Shown are pooled data of three independent experiments with means indicated in red. Statistical significance was assessed by Welch’s ANOVA followed by Dunnett’s T3 multiple-comparison test. *p < 0.05, **p < 0.01. Representative IFA images of the different PVM morphologies are displayed. (F) Total cellular filipin intensity is reduced in OSBP-KD cells. (G) Filipin co-localizes with the PVM marker UIS4. The different masks used for automated image analysis of mean filipin fluorescence intensity are indicated. (H) Filipin enrichment at the parasite relative to the surrounding host cell is reduced in OSBP-KD cells. (I) Pearson’s correlation coefficients between filipin and UIS4 signals within regions including the parasite and adjacent host cell areas are reduced in OSBP-KD cells. For (F), (H) and (I) pooled data of two independent experiments with means indicated in red are shown. Statistical significance was assessed by Welch’s ANOVA followed by Dunnett’s T3 multiple-comparison test. ****p < 0.0001. (J) Proposed model how *Plasmodium* liver stages mimic features of damaged lysosomal membranes at the PVM to co-opt host ER MCS machinery and lipid transfer pathways. Scale bars,10 µm.

To investigate the functional importance of OSBP during liver stage infection, we generated two independent stable OSBP-knockdown (KD) cell lines (OSBP-KD1 and OSBP-KD2) expressing shRNAs targeting OSBP, given the reported essentiality of OSBP^27^. Western blot analysis demonstrated efficient depletion of OSBP expression in both cell lines (Figure 4B). Parasite survival was significantly reduced in both OSBP-KD cell lines, with the stronger phenotype observed in OSBP-KD1 correlating with more efficient depletion of OSBP (Figure 4C). Parasite size at 48 hpi was modestly but significantly increased in both KD cell lines (Figure 4D). In addition, parasites developing in OSBP-KD cells displayed hypervesiculation of the PVM, as quantified by automated image analysis (Figure 4E), indicating that OSBP, similar to VAPA and VAPB, contributes to maintenance of normal PVM morphology.

Because OSBP mediates PI(4)P-dependent cholesterol transport at MCSs^28^, we next examined whether OSBP depletion affects cholesterol distribution during liver stage infection using filipin staining. Analysis of total cellular filipin intensity revealed significantly reduced signals in both OSBP-KD cell lines compared to WT cells (Figure 4F), consistent with the established role of OSBP in cellular cholesterol homeostasis. In line with previous observations that the liver stage PVM is enriched in cholesterol^3^, filipin staining of infected cells revealed partial co-localization with UIS4 at the parasite periphery (Figure 4G). To account for the global reduction in cellular cholesterol, we quantified enrichment at the parasite relative to the surrounding host cell. Using this relative approach, automated image analysis demonstrated significantly reduced cholesterol enrichment at the parasite in both OSBP-KD cell lines (Figure 4H). Quantification of Pearson’s correlation coefficients between filipin and UIS4 signals within regions encompassing the parasite and adjacent host cell areas further confirmed significantly reduced co-localization in both OSBP-KD cells (Figure 4I). Together, these findings suggest that OSBP contributes to cholesterol accumulation at the host-parasite interface and supports maintenance of PVM integrity and parasite survival.

Based on these findings, we propose a model in which *Plasmodium* liver stages establish a PVM that selectively mimics features of damaged endolysosomal membranes (Figure 4J). This membrane state is characterized by PI4K2A-dependent PI(4)P accumulation and recruitment of OSBP to the PVM together with formation of VAPA/B-positive host ER-PVM MCSs. Our data further suggest that OSBP-dependent pathways contribute to cholesterol accumulation at the PVM, while both OSBP- and VAPA/B-dependent pathways support maintenance of PVM integrity and parasite survival.

## DISCUSSION

Our findings establish damage mimicry as a strategy by which *P. berghei* liver stage parasites subvert host membrane repair pathways to support intracellular development. The liver stage PVM recapitulates selected hallmarks of damaged endolysosomal membranes, including sustained CASM responses, PI(4)P accumulation, and recruitment of host lipid transfer machinery, while lacking damage markers such as ESCRT components and galectins. These observations suggest that the parasite selectively engages host membrane stress responses while evading damage recognition pathways that could compromise survival.

A central finding of our study is that the liver stage PVM co-opts a PI4K2A-dependent pathway resembling the phosphoinositide-initiated membrane tethering and lipid transport (PITT) pathway activated during lysosomal membrane repair^16,17^. PI4K2A generates PI(4)P at the PVM and is required for OSBP recruitment, while VAPA/B-positive ER membranes establish MCSs with the PVM. Disruption of either OSBP or VAPA/B impairs PVM integrity and parasite survival, and OSBP depletion reduces cholesterol accumulation at the host-parasite interface. Together, these findings provide a mechanistic framework linking ER-PVM MCSs to lipid transfer and membrane homeostasis during liver stage infection.

The unexpected observation that PI4K2A depletion modestly increased parasite survival, despite reducing PI(4)P accumulation and OSBP recruitment at the PVM, suggests that PI4K2A-associated pathways may originally have evolved as host defense mechanisms engaged at the host-parasite interface, but were subsequently subverted by the parasite to support its intracellular development. Defining how these activities are balanced will require further investigation.

An important open question is whether parasites actively establish the damage-mimicking membrane state. Consistent with this possibility, inhibition of parasite protein export using Torin2 strongly reduced PI4K2A recruitment to the PVM, indicating that exported parasite proteins contribute to this process. Recent studies in *Toxoplasma gondii* demonstrated direct recruitment of VAP proteins by parasite-encoded FFAT motif-containing effectors^29,30^. Our findings point to a distinct mechanism in *Plasmodium*, where establishment of a damage-mimicking membrane identity appears sufficient to recruit the host membrane remodeling machinery, although contributions from parasite-encoded FFAT motif-containing proteins cannot be excluded.

The functional consequence of this damage-mimicking membrane state is likely the activation of host lipid transfer pathways that help meet the extraordinary lipid demands of intracellular replication. Previous studies identified vesicular host cholesterol trafficking pathways that contribute to liver stage development^3,4^. Our findings suggest that non-vesicular cholesterol transfer at ER-PVM MCSs additionally contributes to PVM homeostasis. Consistent with this idea, OSBP depletion reduced cholesterol accumulation at the host-parasite interface and disrupted PVM integrity. Together with the recent identification of the ceramide transfer protein CERT at the PVM^5^, these findings support a model in which multiple host lipid transfer proteins might operate at ER-PVM MCSs.

In conclusion, our findings redefine the liver stage PVM as an engineered metabolic interface that subverts host lysosomal membrane remodeling and lipid transfer pathways to fuel intracellular parasite development. More broadly, they identify damage mimicry as a pathogen strategy for intracellular niche establishment that may extend beyond apicomplexan parasites.

## RESOURCE AVAILABILITY

### Lead Contact

Further information and requests for resources and reagents should be directed to and will be fulfilled by the Lead Contact, Paul-Christian Burda.

### Materials Availability

All plasmids and cell lines generated in this study are available from the Lead Contact upon request.

### Data and Code Availability

All data generated or analyzed during this study are included in this published article and its supplemental material files. All Python-based scripts used for automated image analysis are publicly available and have been archived at Zenodo (https://doi.org/10.5281/zenodo.21011631).

## ACKNOWLEDGEMENTS

We thank Richard J. Youle for providing the HeLa ATG16L1-KO cells. We are grateful to Pietro De Camilli for sharing the VAP-DKO cells and to Harald Stenmark for providing the PI4K2A-KO cell lines. We thank Ross Douglas and Marc Schetelig for providing access to S2 laboratory space and infrastructure at the Technology and Innovation Center Giessen. We are grateful to the MIC (Microscopy Imaging Centre) in Bern for excellent imaging facilities and technical support. This work was supported by the Swiss National Science Foundation (SNSF; grant no. 310030_212795 and CRSII5_198543) to V.T.H. S.H. is supported by a Medical Research Position from the Prof. Dr. Max Cloëtta foundation (Switzerland).

## AUTHOR CONTRIBUTIONS

D.L., J.S.-M., S.H., M.B., R.C., N.U., V.G., A.D.R., A.J.A., P.-C.B. performed the experiments; D.L., J.S.-M., S.H., N.U., P.O., V.T.H., P.-C.B. designed the experiments; D.L., J.S.-M., P.-C.B. prepared the figures; V.T.H. and P.-C.B conceived the project and wrote the paper with input from all authors.

## DECLARATION OF INTERESTS

The authors declare no competing interests.

**Declaration of generative AI and AI-assisted technologies in the writing process** During the preparation of this work the authors used ChatGPT (OpenAI) and Claude (Anthropic) in order to improve the readability and language of this manuscript. After using these tools, the authors reviewed and edited the content as needed and take full responsibility for the content of the published article.

## STAR ★ METHODS

### EXPERIMENTAL MODEL AND SUBJECT DETAILS

#### Animal models

All animal experiments were performed in strict accordance with the guidelines of the Swiss Tierschutzgesetz (TSchG; Animal RightsLaws) and approved by the ethical committee of the University of Bern (license numbers: BE118/2022, BE109/13, BE 81/11). Female and male BALB/c mice used for feeding of mosquitoes with parasites were between 6 and 12 weeks of age and were either purchased at Janvier-Labs (France) or bred in-house. Female and male GFP-LC3 mice^19^ (kindly provided by K. Mizushima) between 6 and 12 weeks of age were used for intravital microscopy and isolation of primary mouse hepatocytes, and were bred in-house. Mice were housed in individually ventilated cages furnished with autoclaved aspen woodchip, a mouse house and paper tissue at 21 ± 2°C under a 12:12 h light-dark cycle at a relative humidity of 55 ± 10%. They were fed a commercially prepared autoclaved dry rodent diet and water, both available *ad libitum*. The health of animals was monitored by routine daily visual health checks. Mice were injected with parasites via an intraperitoneal or intravenous route. The parasitemia of infected animals was determined by flow cytometry. For feeding of mosquitoes, upon reaching a parasitemia of 6%-9% and a gametocytemia of >0.4%, mice were anaesthetized with ketamine (144 mg/kg) and xylazine (19.2 mg/kg), and upon deep anesthesia were placed on a cage with approximately 100 mosquitoes.

#### Mosquito rearing

*Anopheles stephensi* mosquitoes were used for infections with *P. berghei*. They were bred in-house in conditions of 27°C, 80% humidity. Following infection with *P. berghei* parasites, mosquitoes were maintained at 20.5°C at 80% humidity. All mosquitoes were supplied with 8% fructose solution (filter sterilized, supplemented with PABA) and for dissection were anaesthetized in chloroform vapor before submersion in 70% ethanol.

#### Parasite lines

*P. berghei* ANKA parasites were used that constitutively express mCherry in the parasite cytosol^31,32^.

#### Cell lines

HeLa WT cells were obtained from the European Collection of Authenticated Cell Cultures. HEK293T were obtained from the American Type Culture Collection. HeLa cell stably expressing GFP-LC3 were previously generated by our lab^33^. HeLa ATG16L1-KO cells^34^ were kindly provided by Richard J. Youle. PI4K2A-KO cells^16^ were kindly provided by Harald Stenmark. VAP-DKO cells^26^ were a kind gift from Pietro De Camilli. The following cell lines were generated in this study: HeLa WT, PI4K2A-KO and ATG16L1-KO cells stably expressing PI4K2A-HA; HeLa WT, PI4K2A-KO and complemented PI4K2A-KO cells stably expressing the PI(4)P biosensor GFP-ORP9-PH^35^; VAP-DKO cells stably expressing HA-VAPB or HA-VAPBmut; OSBP-KD cells.

### METHOD DETAILS

#### Cell culture and drug treatments

HeLa cells were cultured in Minimum Essential Medium (MEM) with Earle’s salts (BioConcept) or Dulbeccós Modified Eagles Medium (DMEM, Gibco) both supplemented with 10% heat-inactivated fetal bovine serum (Sigma-Aldrich or Gibco), 100 U penicillin, 100 µg/ mL streptomycin (BioConcept or Gibco) and 2 mM L-Glutamine (BioConcept or Gibco) in a humid incubator at 37°C with 5% CO2. Isolation and cultivation of primary mouse hepatocytes was done as previously described^13^. HEK293T cells were cultivated as previously described^33^. Cells were passaged twice a week using trypsin-EDTA (Gibco) or accutase (Innovative Cell Technologies). Cells were treated with DMSO, 20 µM nocodazole (Sigma-Aldrich, M1404), 250 µM LLOMe (Merck, L7393) or 10 nM Torin2 (TargetMol, T6100) as indicated.

#### Molecular cloning

All PCR reactions were performed using Phusion DNA polymerase (NEB). The PI4K2A-HA expression vector pSBbi-Pur-PI4K2A-HA was generated by amplifying the human PI4K2A coding sequence from pPI4K2A-GFP^36^ (kindly provided by Tamas Balla) using primer pair PI4K2A-HA-fw and PI4K2A-HA-rev adding a C-terminal HA-tag. The PCR product was digested with NcoI and HindIII and ligated into the NcoI-and HindIII-digested pSBbi-Pur^37^ (Addgene plasmid #60523). The HA-VAPB expression vectors pSBbi-Pur-VAPB and pSBbi-Pur-VAPBmut were generated by amplifying the human VAPB coding sequence from pEGFP-VAPB^26^ or pEGFP-VAPB (K87D/M89D)^26^ (kindly provided by Pietro De Camilli), respectively using primer pair HA-VAPB-fw and HA-VAPB-rev adding a N-terminal HA-tag. The PCR product was digested with SfiI and ligated into the SfiI-digested pSBbi-Pur. All plasmids were verified by Sanger sequencing.

#### Generation of transgenic host cells

HeLa cells were transfected with 1 to 4 µg of plasmid DNA with program T-28 in an Amaxa Nucleofector 2b (Lonza) using a previously described protocol^38^.

For the generation of cells stably expressing HA-VAPB/HA-VAPBmut and cells stably expressing PI4K2A-HA, the sleeping beauty transposon system^37^ was used. For this, cells were co-transfected with 100 ng of pCMV(CAT)T7-SB100^39^ (Addgene plasmid #34879) and 1.9 µg of each expression plasmid, respectively, followed by puromycin selection as previously described^37^.

For generation of cells stably expressing GFP-ORP9-PH, the plasmid pPBbsr-AcGFP1-ORP9-PH^35^ (Addgene plasmid #214270) was integrated using the piggyBac system^40^ as previously described^35^.

For transient expression of the the microtubule binding-domain of ensconsin fused to mCherry, the plasmid EMTB-mCherry^41^ (Addgene plasmid #26742) was transfected. For transient expression of fluorescent phosphoinositide reporters, the following plasmids were transfected: PI(3)P reporter p40PX-EGFP^42^ (Addgene plasmid #19010), PI(4)P reporter pCMV-AcGFP1-ORP9-PH^35^ (Addgene plasmid #214266), PI(5)P reporter GFP-ING2(pHD)^43^ (Addgene plasmid #21589), PI(3,4)P_2_ reporter GFP-2xPH-TAPP1 WT^44^ (Addgene plasmid #161990), PI(3,5)P_2_ reporter GFP-SnxA (pJSK659)^45^ (Addgene plasmid #205128), PI(4,5)P_2_ reporter GFP-C1-PLCdelta-PH^46^ (Addgene plasmid #21179) and PI(3,4,5)P_3_ reporter pcDNA3-AKT-PH-GFP^47^ (Addgene plasmid #18836). For transient expression of GFP-VAPA and GFP-VAPB, the plasmids GFP-VAPA and GFP-VAPB^48^ (kindly provided by Isabelle Derré) were transfected. For transient expression of GFP-OSBP in WT, PI4K2A-KO and complemented PI4K2A-KO cells, the plasmid pEGFP-OSBP^26^ (kindly provided by Pietro De Camilli) was transfected.

Stable HeLa OSBP-KD cells were generated by lentiviral transduction. Lentiviruses were produced using the lentiviral expression plasmid pLKO.1-OSBP^49^ (Addgene plasmid #32495), the VSV-G envelope plasmid pMD2.G (Addgene plasmid #12259), and the second-generation packaging plasmid psPAX2 (Addgene plasmid #12260). pMD2.G and psPAX2 were gifts from Didier Trono. Lentivirus production and transduction were performed as previously described^33^.

#### Immunoblotting

Protein samples were resolved by SDS-PAGE and transferred to nitrocellulose membranes (LI-COR). Membranes were blocked in 5% milk in TBS-T, which was also used for subsequent antibody incubations. The following primary antibodies were used: rabbit anti-OSBP (Proteintech, 11096-1-AP; 1:1,000) and chicken anti-GAPDH (Millipore, AB2302; 1:5,000). The following secondary antibodies were used: goat anti-rabbit IgG IRDye 800CW (LI-COR, 926-32211; 1:10,000) and donkey anti-chicken IgG IRDye 680LT (LI-COR, 926-32218; 1:10,000). Proteins were detected using an Odyssey Imaging System (LI-COR). Quantifications were performed using ImageJ^50^.

#### Liver stage infections

HeLa cells were seeded in a 96-well plate at 40,000 cells/well the day before infection and were infected with ∼20,000 *P. berghei* sporozoites for 2 h. The infected cells were then detached using accutase and seeded at lower density. For certain experiments, cells were already plated at lower density on coverslips or live-cell imaging dishes and directly infected without re-seeding. For infection of cells, salivary glands of infected *A. stephensi* mosquitoes were isolated and disrupted to release sporozoites. Sporozoites were incubated with cells in medium containing 2.5 μg/mL amphotericin B (Bioconcept, Biowest) to prevent fungal contamination. Medium was changed 2 hpi and then once per day.

#### Liver stage analysis

Automated live-cell imaging was used to determine parasite size and numbers. Infected cells were seeded in a 96-well format and imaged using a 10x objective with an INCell Analyzer 2000 automated live cell imaging system (GE Healthcare Life Sciences) at 6 hpi and 48 hpi with minimal light exposure. INCell Developer Toolbox 1.10.0 software was used to analyze the acquired images. The mCherry cytoplasmic signal was used to determine the parasite area. Image analysis parameters (kernel size, sensitivity, and size threshold) were optimized to accurately segment parasites from background. Survival rates 6 to 48 hpi were normalized to 6 hpi parasite counts (set as 100% viability baseline). Parasite area (size) was assessed at 48 hpi.

#### Immunofluoresence analysis

At different timepoints post-infection, cells were fixed using 4% paraformaldehyde (PFA) in PBS for 10 to 20 min at room temperature (RT). Permeabilization was performed for 10 min in 0.1 - 0.25 % Triton X-100 in PBS or in icecold methanol. After washing with PBS, unspecific binding sites were blocked by incubation in 10% FCS/PBS for 10 to 30 min followed by incubation with primary antibody in 10% FCS/PBS for 1 h. After washing with PBS, cells were incubated with fluorescently labeled secondary antibodies diluted in 10% FCS/PBS for 1 h. DNA was stained with 1 µg/mL DAPI during secondary antibody incubation. After washing another time in PBS, cover slips were mounted onto glass slides using ProLongTM Gold antifade solution (Invitrogen, P36930) or ProLong Diamond (Molecular Probes, 15372192). The following primary antibodies were used: rat monoclonal anti-red antibody (Chromotek, 5f8; 1:1,000), mouse monoclonal anti-GFP antibody (Roche, 11814460001; 1:1,000), mouse monoclonal anti-mNeonGreen (Chromotek, 32F6; 1:1,000), rabbit anti-UIS4 antiserum (Proteogenix; 1:1,000 - 10,000) and rat anti-HA high affinity (Roche, #45-11867423001; 1:1,000). The following secondary antibodies were used: Goat anti-Rat IgG (H+L) Highly Cross-Adsorbed Secondary Antibody, Alexa Fluor™ Plus 594 (Invitrogen, A48264; 1:2,000), Goat anti-Rabbit IgG (H+L) Cross-Adsorbed Secondary Antibody, Alexa Fluor™ 488 (Invitrogen, A-11008; 1:2,000), Goat anti-Rat IgG (H+L) Cross-Adsorbed Secondary Antibody, Alexa Fluor™ 488 (Invitrogen, A11006; 1:1,000), Goat anti-Mouse IgG (H+L) Cross-Adsorbed Secondary Antibody, Alexa Fluor™ 488 (Invitrogen, A-11001; 1:2,000), Goat anti-Mouse IgG (H+L) Highly Cross-Adsorbed Secondary Antibody, Alexa Fluor™ 488 (Thermo, A-11029; 1:1,000), goat anti-rabbit CY5 (Jackson Immuno Research, 111-175-144; 1:1,000) and goat anti-Rabbit StarRed (Abberior, STRED-1002; 1:2,000).

#### Proximity ligation assay

The proximity ligation assay was performed using the Duolink In Situ reagents (Sigma-Aldrich) basically according to manufacturer’s instructions. Infected cells in optical 96-well plates were fixed at 24 hpi with 4 % PFA in PBS for 10 min and permeabilized with 0.1 % Triton X-100 in PBS for 10 min at RT. After blocking, cells were incubated for 30 min with rabbit anti-UIS4 (Proteogenix, 1:20,000) and mouse anti-HA (Santa Cruz, SC-7392, 1:1,000) antibodies at RT. Incubation with PLA probes anti-rabbit PLUS (Sigma-Aldrich, DUO92002) and anti-mouse MINUS (Sigma-Aldrich, DUO92004), washing steps (Sigma-Aldrich, DUO82049) as well as ligation and amplification (Sigma-Aldrich, DUO92014) were performed according to the manufacturer’s instructions. Following amplification, cells were incubated with goat anti-rabbit StarRed secondary antibodies (Abberior, STRED-1002; 1:2,000) for 30 min at RT to visualize UIS4 and with DAPI (1 µg/mL) to stain nuclei. Cells were kept in PBS until imaging.

#### Filipin staining

Infected cells on coverslips were fixed at 30 hpi with 4% PFA in PBS for 10 min (all incubations at RT). Following two washes with PBS, cells were stained with 150 μg/mL Filipin III (MedChemExpress, HY-N6718; prepared from a 5 mg/mL stock in DMSO) in PBS for 1 h. Cells were then washed twice with PBS before permeabilization with 0.1 % Triton X-100 for 10 min. After washing three times with PBS, cells were blocked with 10% FCS in PBS for 10 min and incubated with chicken anti-UIS4 antibody (Proteogenix; 1:5,000 in 10% FCS/PBS) for 1 h. Cells were washed twice with PBS and incubated with highly cross-adsorbed donkey anti-chicken secondary antibody labelled with Alexa Fluor 488 (Invitrogen, A78948; 1:1,000) in 10% FCS/PBS. Following two final washes with PBS, coverslips were mounted on glass slides using ProLong Gold antifade solution (Invitrogen, P36930) and imaged on the same day.

#### Microscopy

Live-cell imaging and time-lapse microscopy of infected GFP-LC3-expressing cells was performed using a Leica DMI6000B widefield epifluorescence microscope equipped with a Leica HCX Plan-Apochromat CS 63x/1.2 water immersion objective. Live-cell imaging of fluorescent phosphoinositide reporters was done using a Nikon Ti2 microscope equipped with Crest V-light V3 spinning disc confocal system with a CFI Plan-Apochromat Lambda 100X /1.45 oil objective. During imaging, cells were kept in 5 % CO_2_ at 37°C. Intravital imaging was performed as previously described ^51^ using a confocal LSM 510 Zeiss microscope equipped with a Zeiss Plan-Apochromat 63x/1.4 oil DIC M27 objective. PLA samples were imaged with an inverted Leica TCS SP8 confocal microscope using an HC Plan-Apochromat CS2 63x/1.4 oil objective. IFA samples were imaged using the Leica TCS SP8 confocal microscope or a Zeiss Axio Observer.Z1 inverted widefield microscope equipped with an ApoTome.2 structured illumination module for optical sectioning with a Plan-Apochromat 63x/1.4 oil objective. Filipin-stained samples were imaged using a Leica DM5500B widefield microscope with a HCX Plan-Apochromat 100x/1.4 oil objective. Image processing was performed using ImageJ^50^.

### QUANTIFICATION AND STATISTICAL ANALYSIS

#### Analysis of PI4K2A and PI(4)P localization

Quantification of PI4K2A-HA-positive parasites in WT and ATG16L1-KO cells was done by visual inspection using a Leica DM5500B epifluorescence microscope with a 63x objective. Only UIS4-positive parasites were considered. Parasites were counted positive if a clear association between PI4K2A-HA and UIS4 at the parasite circumference was observed which means that more than 30% of the proximity of the parasite was covered with the protein of interest. The same strategy was performed for quantification of PI(4)P-positive parasites in WT, PI4K2A-KO and complemented PI4K2A-KO cells. Quantification of relative PI(4)P reporter intensity in these cells was determined using ImageJ^50^.

#### Automated image analysis

Automated image analysis pipelines were implemented in Python with assistance from Claude (Anthropic). For each workflow, analysis outputs, including segmentation masks and object classifications, were visually inspected on experimental images to verify accurate identification of the biological structures used for quantification. Within each experiment, all samples were imaged using identical acquisition settings.

#### Analysis of Torin2 treated parasites

For determination of PI4K2A-HA and UIS4 localization of parasites in DMSO- and Torin2-treated HeLa cells, single parasite images were cropped to 27 x 27 µm. The parasite body was segmented from the mCherry channel by Gaussian smoothing (σ = 0.18 µm), Otsu thresholding at 50% of the computed threshold value, morphological closing (disk radius 0.54 µm), hole filling, and removal of objects smaller than 2.6 µm². The component closest to the image center was retained and subsequently eroded by 0.36 µm to exclude PVM-proximal signal from the parasite interior. Concentric compartments were defined relative to the eroded mask boundary using a Euclidean distance transform: a PVM ring (0-1.07 µm outside the mask) and a host-cell reference annulus (1.61-7.05 µm outside the mask, separated from the PVM ring by a 0.54 µm gap). Mean UIS4 and PI4K2A fluorescence intensity were measured in each compartment. UIS4 export was expressed as the ratio of mean UIS4 intensity in the PVM ring to mean UIS4 intensity within the parasite mask. PI4K2A enrichment at the PVM was expressed as the ratio of mean PVM ring intensity to mean host annulus intensity.

#### Quantification of PVM-associated PLA dots

For quantification of PVM-associated PLA dots, the PVM mask was generated by Otsu thresholding of the PVM marker channel after Gaussian smoothing (3×3-pixel kernel); connected components smaller than 0.17 µm² were discarded as noise. PLA dots were detected by applying a Difference-of-Gaussians (DoG) filter (σ₁ = 0.091 µm, σ₂ = 0.228 µm) to the PLA channel, followed by local-maximum detection requiring a minimum center-to-center separation of 0.273 µm and a DoG-filter amplitude of at least 5% of that image’s maximum DoG response (with a floor of one detector count for signal-free images); candidate maxima were then required to exceed a minimum raw-intensity threshold of 25 detector counts to be retained as PLA dots. Each retained PLA dot was classified as PVM-proximal or cytosolic by computing the Euclidean distance from the dot center to the nearest pixel of the PVM mask; dots within 0.25 µm of the PVM mask were classified as PVM-proximal, and dots beyond this threshold were classified as cytosolic and excluded from further analysis.

#### Quantification of PVM vesiculation

For quantification of PVM vesiculation, images were cropped to 50 × 50 µm prior to analysis. For segmentation, the UIS4 channel was contrast-normalized by percentile stretching (1st–99.5th percentile) to an 8-bit range, followed by Gaussian blurring (3×3 kernel, σ = 0.5 px). Binary segmentation was performed using an Otsu threshold scaled by a factor of 1.2 to retain bright PVM signal while suppressing dim background. Morphological closing with an elliptical structuring element (4-pixel diameter) was applied to connect membrane gaps. Connected-component analysis (8-connectivity) was then performed on the binary mask. Each connected component with an area ≥ 0.05 µm² was retained for further classification. A mCherry mask was generated independently by Otsu thresholding of the mCherry channel after Gaussian blurring (3×3 kernel, σ = 0.5 px), morphological closing, and hole filling, without further dilation. The largest UIS4-positive component overlapping the mCherry-positive parasite body was identified as the intact PVM main body and always counted as one structure. Components whose centroid fell within the segmented mCherry mask were excluded as internal signal rather than counted as additional detached vesicles. All remaining components were counted as detached UIS4-positive PVM vesicles. The total number of PVM vesicles per parasite equals the number of detached vesicles plus one.

#### Quantification of GFP-OSBP localization

For co-localization analysis between OSBP-GFP and the PVM marker UIS4, images were cropped to 40 x 40 µm. The GFP and UIS4 channels were extracted and analyzed without spatial masking or background subtraction. Co-localization was expressed as the Pearson correlation coefficient computed over all pixels.

#### Quantification of filipin staining

For measurement of filipin fluorescence intensity, two complementary metrics were measured. First, filipin staining intensities in total cells (infected and uninfected) were assessed from uncropped field-of-view images. The filipin channel was smoothed with a Gaussian filter (σ = 0.18 µm), and the cellular foreground was segmented by thresholding at the intensity peak of the darkest 5% of pixels plus a fixed offset of 40 grey-level units. Connected components smaller than 50 µm² were discarded as debris, and residual holes within foreground regions smaller than 50 µm² were filled. Mean filipin intensity was reported for foreground pixels of the unsmoothed image. Second, cholesterol enrichment at the parasite was quantified from 26 × 26 µm cropped images with single parasites. The parasite was segmented using the UIS4 channel by Otsu thresholding after Gaussian smoothing (σ = 0.18 µm), followed by morphological closing (radius 0.54 µm), hole filling, and removal of objects smaller than 5 µm². A local background annulus was defined as the region between 0.27 µm and 2.28 µm outside the parasite mask boundary, calculated via Euclidean distance transform. The parasite/background ratio was computed as the mean filipin intensity within the parasite mask divided by the mean filipin intensity in the background annulus. Colocalization between UIS4 and filipin was quantified as the Pearson correlation coefficient within a region of interest dilated by 1.37 µm beyond the parasite mask boundary.

### Statistical analysis

Comparisons between two independent groups were performed using Welch’s t-test. Comparisons among multiple independent groups were performed using Welch’s ANOVA followed by Dunnett’s T3 multiple-comparison test. Complete matched datasets were analyzed using repeated-measures one-way ANOVA followed by Holm–Šidák multiple-comparison testing. Matched datasets in which not all treatment groups were available in every independent experiment were analyzed using a mixed-effects model (REML) with experiment treated as a matched factor, followed by Holm–Šidák multiple-comparison testing. All statistical tests were performed in GraphPad Prism 11. P values <0.05 were considered statistically significant. Statistical details, including the number of independent experiments, statistical tests used, and definition of error bars, are provided in the figure legends.

## SUPPLEMENTAL MOVIES

**Movie S1. Formation of dynamic LC3-positive PVM tubules *in vitro*. Related to Figure 1**. Live-cell time-lapse microscopy of *P. berghei*-infected host cell expressing GFP-LC3 at 4 hpi. Images were acquired at 1 sec intervals. Scale bar, 10 µm.

**Movie S2. Formation of LC3-positive PVM tubules *in vivo.* Related to Figure 1**. Intravital microscopy movie of a *P. berghei*-infected GFP-LC3 mouse at 4 hpi. The parasite is shown in magenta, while the GFP-LC3 signal is displayed in green. Images were acquired at 4 sec intervals. Scale bar, 10 µm.

## Notes

https://doi.org/10.5281/zenodo.21011631

### Competing Interest Statement

The authors have declared no competing interest.

