## Supplemental Information for "*Plasmodium berghei* liver stages establish a damage-mimicking vacuole to hijack host ER membrane contact site machinery for lipid uptake"

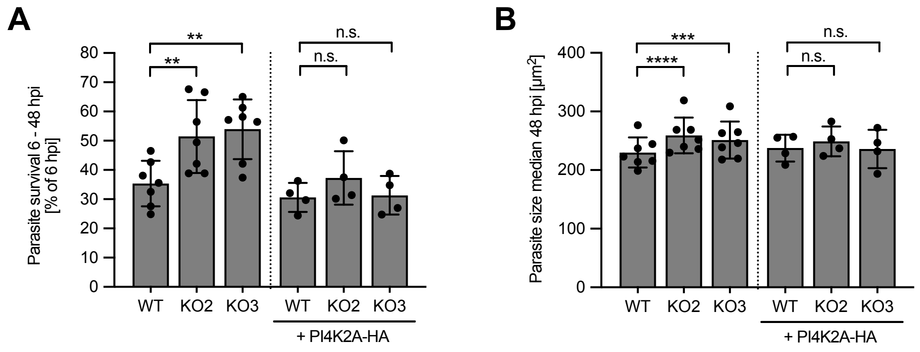


**Figure S1. Knockout of host PI4K2A does not impair parasite development. Related to Figure 2.** (A) Parasite survival (6 to 48 hpi) in two independent PI4K2A-KO cell lines is increased in comparison to WT cells. This effect is restored upon PI4K2A complementation. (B) Parasite size at 48 hpi is slightly increased in PI4K2A-KO cell lines and this effect is restored upon PI4K2A complementation. Shown are means ± SD of four to six independent experiments. Statistical significance was assessed by repeated-measures one-way ANOVA followed by Holm–Šidák multiple-comparison testing. $^{**}p < 0.01$, $^{***}p < 0.001$, $^{****}p < 0.0001$; n.s., not significant.

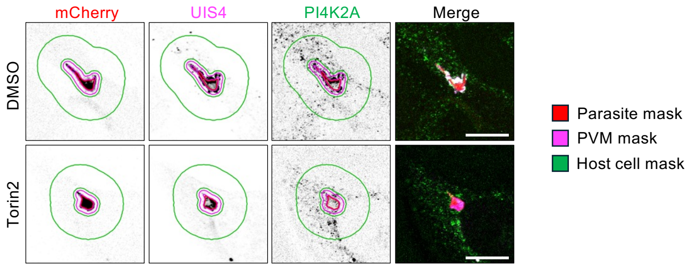


**Figure S2. Automated image analysis of PI4K2A recruitment upon Torin2 treatment. Related to Figure 2.** Parasite, PVM and host cell masks used for Python-script-based image analysis are shown for a representative DMSO- and Torin2-treated parasite. Scale bars, 10 µm.
